# SUMOylation of GMFB regulates the stability and function of GMFB in RPE cells under oxidative stress and inflammation

**DOI:** 10.1101/2021.03.03.433763

**Authors:** Wan Sun, Juan Wang, Jieping Zhang, Furong Gao, Qingjian Ou, Haibin Tian, Caixia Jin, Jingying Xu, Jingfa Zhang, Jian huang, Guo-Tong Xu, Lixia Lu

## Abstract

Glia maturation factor beta (GMFB) is a growth and differentiation factor that act as an intracellular regulator of signal transduction pathways. The SUMOylation is a post-translational modification (PTM) that plays a key role in protein subcellular localization, stability, transcription, and enzymatic activity. Recent studies have highlighted the importance of SUMOylation in the inflammation and progression of numerous diseases. But little is known about the relationship between GMFB and SUMOylation. Here we first report that GMFB can be mono-SUMOylated at multiple sites by the covalent addition of a single SUMO1 protein, and identified K20, K35, K58, and K97 as major SUMO acceptor sites. We also found that SUMOylation leading to increased stability and trans-localization of GMFB. Furthermore, RNA-seq data and Real-time quantitative polymerase chain reaction (rt-qPCR) also indicated that the SUMOylated GMFB upregulated multiple pathways, including the cytokine-cytokin receptor interaction, NOD-like receptor signaling pathway, TNF signaling pathway, RIG-I-like receptor signaling pathway, and NF-kappa B signaling pathway. Our studies intend to provide a novel direction for the study into the biofunction of GMFB, SUMOylated GMFB and the mechanism, clinical therapy, and prognosis of inflammation-related RPE disorders like age-related macular degeneration (AMD) and diabetic retinopathy (DR).

## Introduction

Glia maturation factor-β (GMFB), a highly evolutionary conserved protein, is a 17-kDa growth and differentiation factor that contains an ADF-H domain (1,2). GMFB protein is primarily expressed in the brain and also present in the inner layer of the retina, a special part of the CNS(3,4). GMFB involved in Parkinson’s disease, breast cancer, neuroinflammatory conditions including Multiple Sclerosis and so on(5–7). Over-expressed GMFB was more frequently localized in the microvascular endothelial cells in tumor cells in high-grade and co-localized with CD31, which means that GMFB probably related to the formation of Neovascularization(8). GMFB participates in many inflammatory and immune reactions. The phosphorylated GMFB is an enhancer of p38 mitogen-activated protein kinase (MAPK) and an inhibitor of extracellular signal-regulated kinase 1 and 2 (ERK1/ERK2)(9,10).

Post-translational reversible modification by small ubiquitin-related modifiers (SUMO) is known to regulate protein stability, localization, and activity(11). There are four SUMO isoforms that have been identified, SUMO1, SUMO2/3 and SUMO4, which mediating SUMOylation in mammals(12,13). Being a member of the SUMO family, SUMO1 participates in growth, development, and disease in mammals(14–16). SUMOylation plays fundamental roles during eye development and pathogenesis and involved in multiple cellular processes, such as regulation of transcription, apoptosis, transport to the nucleus, protein stability, cellular stress response and cell cycle progression (17–19). Recently an increasing number of studies have connected SUMOylation to the pathological changes of eye diseases, including diabetic retinopathy, UVB-induced retinal diseases, retinal dystrophies and so on(20–22). The article by Hu et al. demonstrated that in rat and cellular model systems SUMOylation regulates Nox1-mediated DR by inhibiting ROS generation and apoptosis(23). SUMOylation also regulates NF‐κB signaling in glomerular cells from diabetic rats and is associated with pathological angiogenesis through regulates the Vascular endothelial cell growth factors receptor 2 (VEGFR2) by SUMOylation and deSUMOylation (24,25).

We note that GMFB and SUMOylation are associated with the regulation of inflammation. Also, it is well known that retinal pigment epithelial (RPE) cells play crucial roles in several ocular pathologies. RPE cell damage is the hallmark of AMD pathogenesis, and one of the mechanisms is oxidative stress and inflammation(26). Dysfunction of RPE causes the imbalance of secretions of cytokines, chemokines, and growth factors in DR, which are related to inflammation, reduction permeability, excessive production of reactive oxygen species (ROS), and apoptosis (27,28).

To date, there are no published studies on the biofunction of SUMOylated GMFB. Therefore, we hypothesized that GMFB may be modulated by SUMOylation and play essential roles in RPE cells under oxidative stress and inflammation. We detect the expression pattern of GMFB and SUMO in retina tissues, and whether SUMOylation modulates the stability and function of GMFB. We also used RNA-seq to investigate the function of SUMOylated GMFB. Here, our findings reveal that GMFB can be mono-SUMOylated at multiple sites. Furthermore, GMFB and SUMOylated GMFB correlates closely with oxidative stress and inflammation.

## Results

### GMFB and protein SUMOylation enhances in RPE cells at early stages of oxidative stress and inflammation

First, we aimed to characterize the expression patterns of GMFB at the early stages of oxidative stress and inflammation. The blood glucose level and body weight changes were detected after Streptozotocin (STZ) or saline injection in Sprague Dawley rats to ensure that modeling was successful (Fig. 1A, B). The expression of GMFB was detected in the total retina of the STZ-induced diabetic rats by rt-qPCR and western blot analysis. Significantly increased expression of GMFB was observed in 1W and 2W, and then decreased at 7 weeks compared to the control (Fig. 1C). The results of the immunofluorescence assay reveal that GMFB is expressed in multiple cell types including RPE cells (Fig. 1D). Moreover, we detect the expression patterns of SUMO1, SUMO2 and SUMO3 in neural retina and RPE-Bruch's membrane-choriocapillaris complex (RBCC). The expression patterns of SUMOs in the neural retina were the same as in RBCC. SUMO2 shows the highest expression level followed by SUMO1, and SUMO3 had the lowest level of expression (Fig. 1E). We also detected the expression patterns of SUMOs in RBCC under oxidative stress and inflammation in the sodium iodate-inducible model(29). The result indicated that the expression of SUMO1 and SUMO3 significantly increased first and then temporarily decreased and increase again under oxidative stress and inflammation, and the expression level of SUMO1 is much higher than SUMO3(Fig 1F). There is a low expression in 12h, because of apoptosis of RPE cell observed at 12 hours post-injection in sodium iodate-inducible model (30). Then we detect the expression and location of GMFB and SUMO1 in ARPE-19 cells with or without high glucose treatment by immunofluorescence, and found that high glucose treatment could significantly promote GMFB/SUMO1 co-localization (Fig 1G). These results showed that SUMO1 and GMFB were both expressed in RPE cells, and the expression both increased under the early stage of oxidative stress and inflammation.

**Figure 1.**
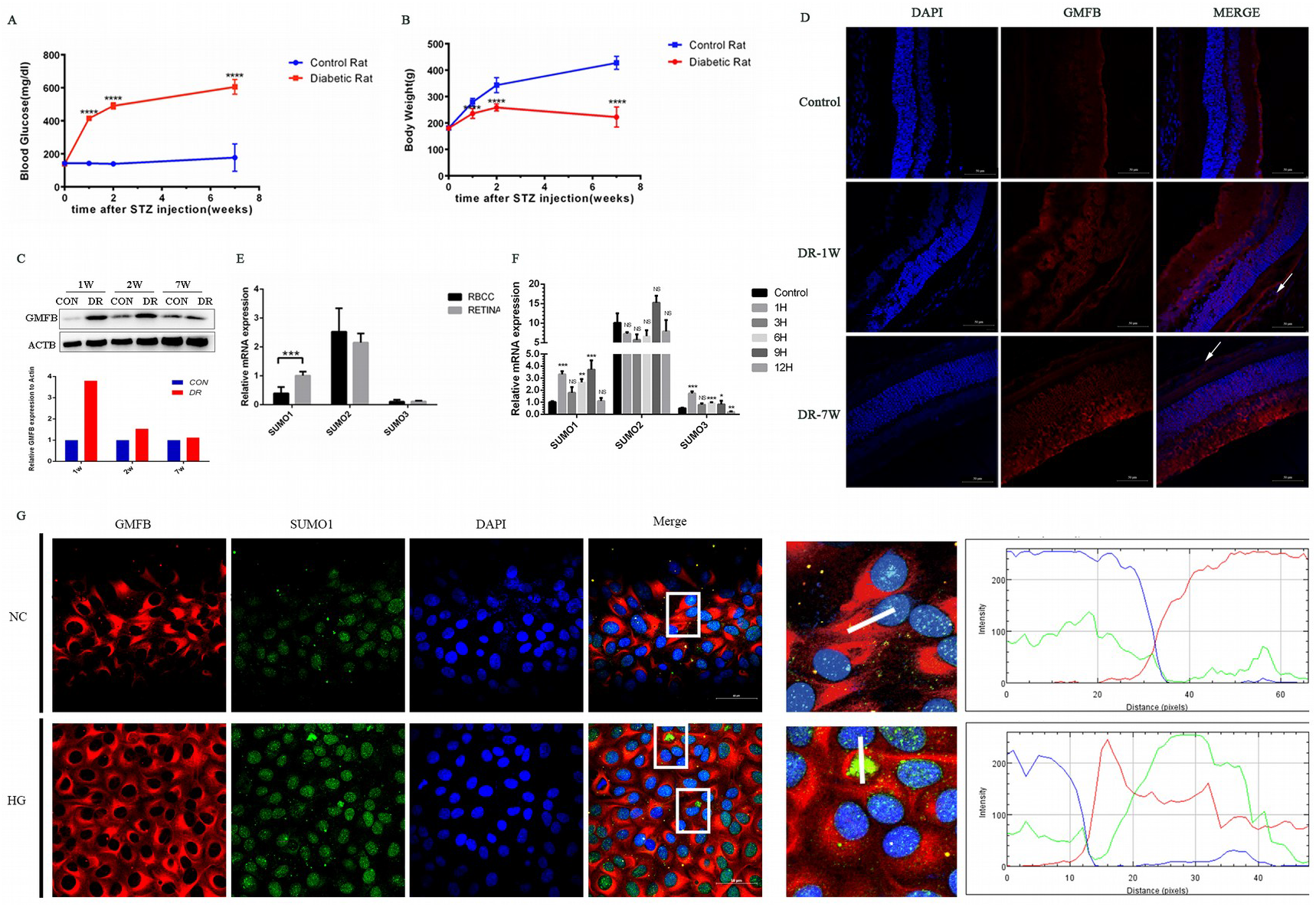
GMFB and protein SUMOylation enhances in RPE cells at early stages of oxidative stress and inflammation. Blood glucose (A) and body weight (B) of normal and STZ-induced diabetic rat in first 7 weeks after the STZ treatment; (C) GMFB expression level in RBCC OF STZ-induced diabetic rat at 1 week, 2 week and 7 weeks were determined by western blotting, and gray analysis of western blotting; (D) The retina of control, 1 week and 7 weeks DR rat were stained by immunofluorescence with anti-GMFB antibody and DAPI, white arrows: RPE layer; Scale bar represents 50 µm. (E) Relative expression of SUMO1, SUMO2 and SUMO3 in neural retina and RBCC; (F) Relative expression of SUMO1, SUMO2 and SUMO3 in the sodium iodate-inducible model of oxidative injury to the RPE; Bars show the relative expression level of mRNA for each condition according to the comparative Ct method: Ratio = 2-ΔΔCt. * p<0.05, ** p<0.001, *** p<0.0001, nonspecific (NS). (G) Immunofluorescence of GMFB and SUMO1 in ARPE-19 cells treated with high glucose for 24 hours. Channel intensity profiles for the red and green channels were performed by Image J. Scale bar represents 50 μm.

### GMFB is modified by SUMO1, and the SUMOylated site is shifted

As presented in Fig. 2A, SDS-PAGE analysis revealed that a predicted band at about 38KD appeared with anti-GMFB, when co-transfected with FLAG-GMFB and His-SUMO1. And the nuclear 38 kDa GMFB protein was gradually increased with UBC9 (SUMO E2 ligase), and significantly reduced with SENP1(deSUMO conjugase). It was demonstrated that the protein band at about 38KD is a GMFB-SUMO1 specific band, which means there is only one “K” on GMFB protein can be mono-SUMOylated by SUMO1. We also noticed that when co-transfected SUMO1 with GMFB, the expression level of GMFB is higher, which indicated that SUMOylation may enhance the stability of GMFB. Most SUMO-modified proteins contain the consensus motif, ψ-K-x-D/E, where ψ is a hydrophobic residue, K is the lysine conjugated to SUMO, x is any residue and D/E is an acidic residue. To determine which lysine of GMFB is potential SUMO modification sites, we performed predictions of sumoylation sites based on a direct amino acid match to SUMO-CS and substitution of the consensus amino acid residues with amino acid residues exhibiting similar hydrophobicity through the UbPred (http://www.ubpred.org/), SUMOplot (http://www.abgent.com/sumoplot/), and GPS-SUMO(http://sumosp.biocuckoo.org/online.php)(31). There are three predicted SUMOylation sites, 35K, 58K and 137K, and two SUMO-interaction motifs located at residues 5-9 (SIM5-9) and residues 42-46 (SIM42-46) on GMFB (Table 1, Fig. 2B). Multiple sequence alignment showed that all residues K in GMFB protein are highly conserved across species (Fig. 2B). First, we selected three predicted SUMOylated sites for experimental validation. However, the specific SUMOylated site of GMFB was not found (Fig.2B-C). Thus, we constructed Flag-GMFB K to R series expression constructs, co-expressed with or without His-SUMO1 and UBC9 in HEK293T(Fig.2C). And the result indicated that almost all mutant K to R GMFB proteins still can be SUMOylated by SUMO1(Fig.2D-F). These data indicated that although GMFB can be SUMOylated at only one “K” site, there is no specific SUMOylated site on GMFB. The SUMOylated “K” site is shifted.

**Figure 2.**
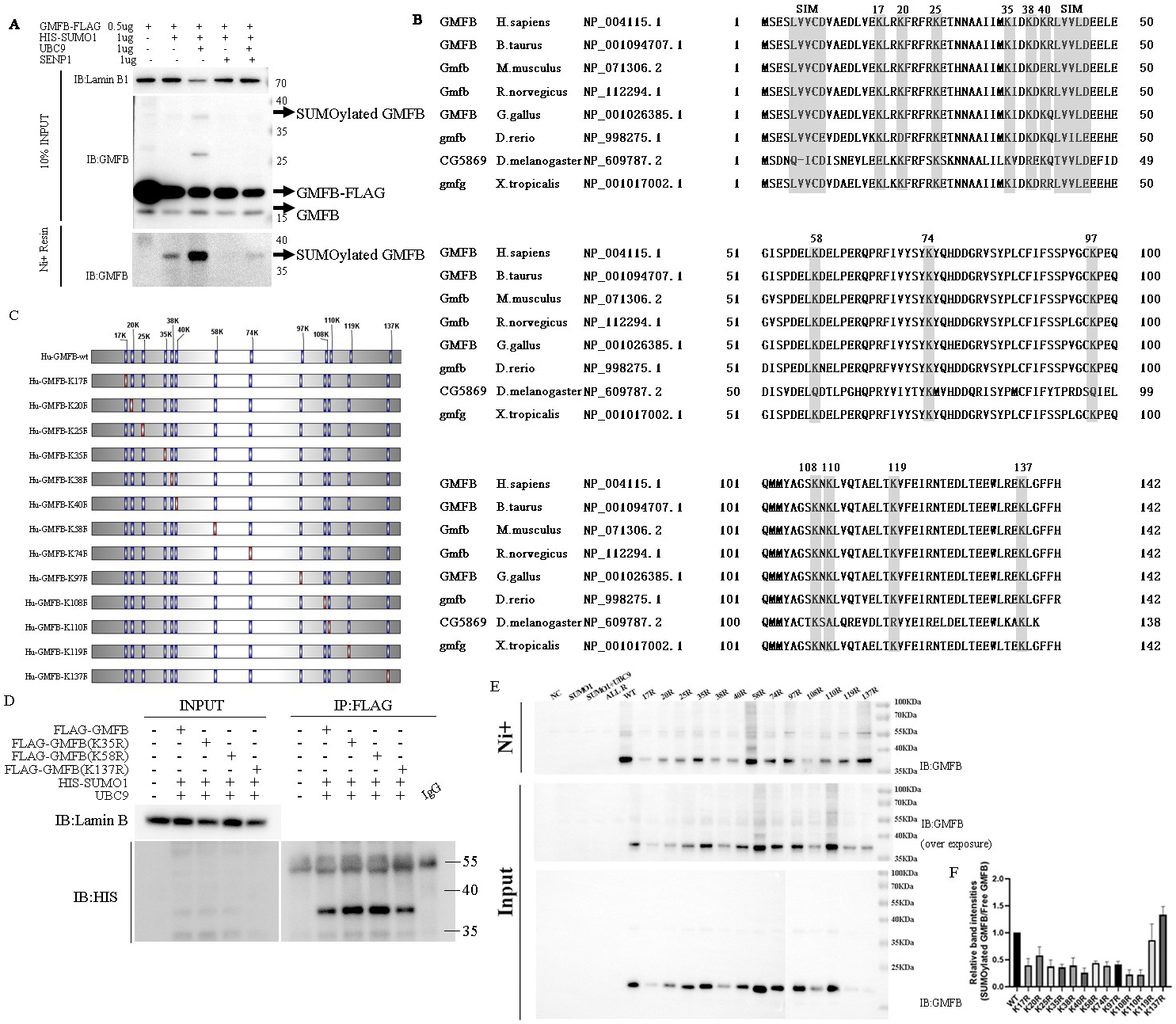
GMFB is modified by SUMO1, and the SUMOylated site is shifted (A) 293T cells were transfected with expression vectors for GMFB-WT-Flag(0.5ug), His-SUMO1(1ug), UBC9(1ug), and SENP1(1ug). And the protein extracts were immunoblot with antibody to GMFB. (B) Conservation of amino acids in predict post-translational modification sites. The gray areas indicate the sites which predicted post-translational modification. (C) Schematic of GMFB K to R constructs. (D) 293 cells co-transfected with His-SUMO1 and Flag-GMFB(WT), Flag-GMFB (K35R), Flag-GMFB (K58R) and Flag-GMFB (K137R) were IP for FLAG followed by IB for His (SUMO1). (E) 293 cells co-transfected with His-SUMO1and Flag-GMFB(WT), Flag-GMFB (K to R) series plasmids were IP for His(SUMO1) followed by IB for GMFB. (G)The ratio of SUMOylated GMFB to the free GMFB of the quantitative analysis of the gray density of Western blotting bands.

**Table 1.** Prediction of SUMOylation Sites in GMFB.

| Human GMFB | UbPred | SUMOplot | GPS-SUMO |
| --- | --- | --- | --- |
| K17 | - | - | - |
| K20 | - | - | - |
| K25 | - | - | - |
| K35 | - | + | + |
| K38 | - | - | - |
| K40 | - | - | - |
| K58 | + | + | + |
| K74 | - | - | - |
| K97 | - | - | - |
| K108 | - | - | - |
| K110 | - | - | - |
| K119 | - | - | - |
| K137 | - | + | + |

### K20, K35, K58 and K97 are the major SUMOylation site of GMFB

To confirm that GMFB is SUMOylated at alternative sites, we constructed HA-GMFB K only series expression constructs, which mutants with only one lysine on GMFB protein. We first detect the correct expression of our K-only series plasmid, then co-transfected with His-SUMO1 and UBC9 to detected the SUMOylated GMFB band. Our result showed that GMFB can be SUMOylated at K20, K25, K35, K58, K74, K97, K108, K110, K119, K137, but not K17, K38 and K40 (Fig. 3C, D). And based on relative band intensities of SUMOylated GMFB, we indicated K20, K35, K58 and K97 are the major SUMOylation site of GMFB.

**Figure 3.**
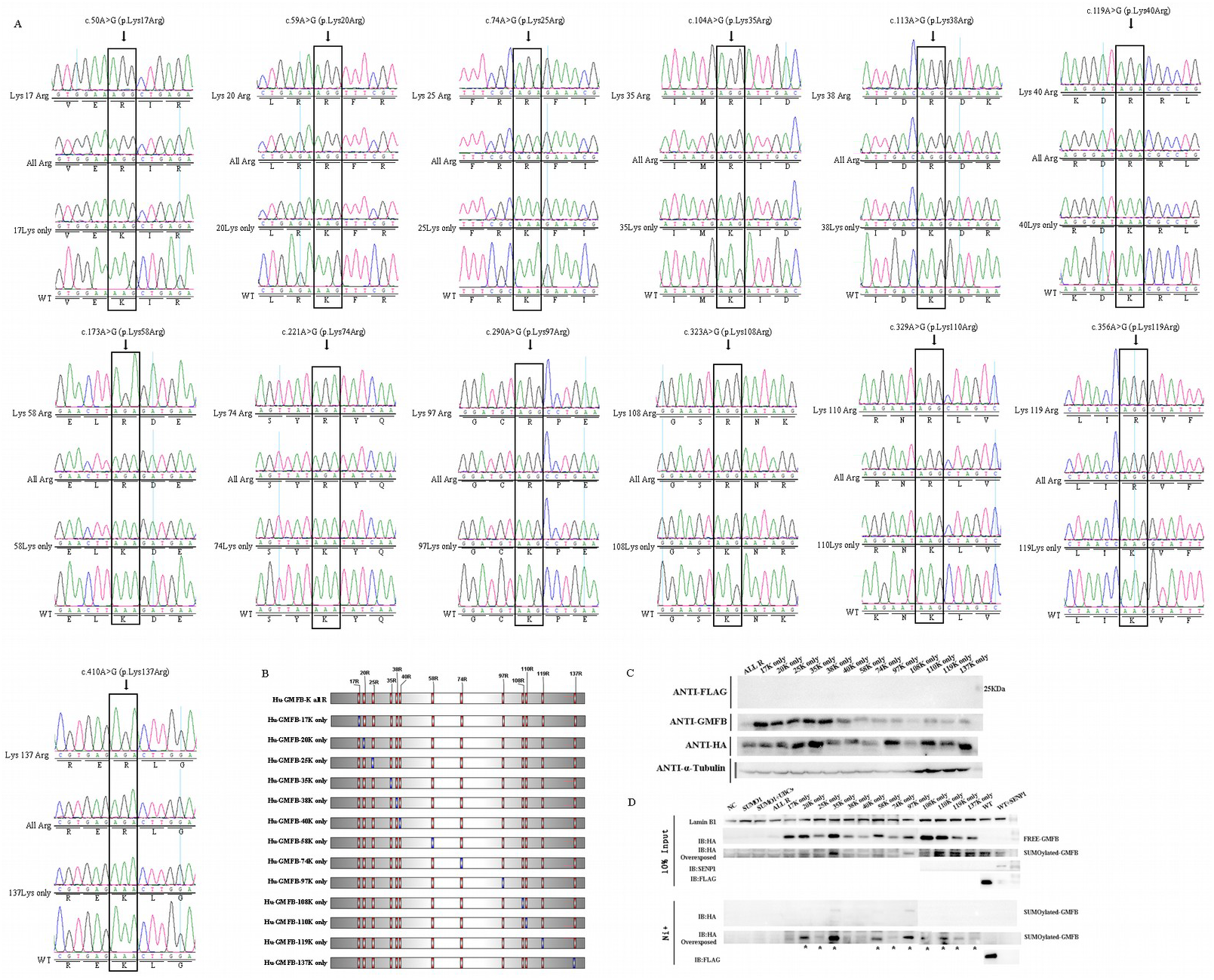
K35, K58 and K97 are the major SUMOylation site of GMFB (A) Partial sequence electropherograms of GMFB mutants K-all-R and GMFB-17K-ONLY, GMFB-20K-ONLY, GMFB-25K-ONLY, GMFB-35K-ONLY, GMFB-38K-ONLY, GMFB-40K-ONLY, GMFB-58K-ONLY, GMFB-74K-ONLY, GMFB-97K-ONLY, GMFB-108K-ONLY, GMFB-110K-ONLY, GMFB-119K-ONLY, GMFB-137K-ONLY plasmid and the correspond GMFB-K to R plasmid. (B) Schematic of GMFB constructs. (C) 293 cells were transfected with plasmids for expression of the K-only GMFB protein. Protein levels were analyzed by Western blotting showing that K-only GMFB plasmids were well expressed in 293 cells. We generated a rabbit polyclonal anti-GMFB antibody to indicate the sensitivity for the detection of the mutant protein. (D) 293 cells were transfected with His-SUMO1 and HA-GMFB (K only) plasmids were IP (HA) followed by IB for GMFB.

### GMFB SUMOylation regulates protein stability

Protein regulation by reversible attachment of SUMO plays an important role in protein stability. To detect whether there are differences in stability between the GMFB wt and GMFB all R, total cell lysates from 293 cells transfected with GMFB wt-FLAG or GMFB all R-FLAG post-treated with DMSO or CHX were measured. Samples from DMSO and CHX treated cells were analyzed separately to prevent incorrect imputation. The result revealed that GMFB wt protein is very stable, but GMFB all R is not (Fig. 4 A-C). Aim to know whether GMFB degradation is mainly dependent on the proteasomal or lysosomal pathway, we measured the protein level in total cell lysates from 293 cells transfected with GMFB wt-FLAG or GMFB all R-FLAG post-treated with MG132 or chloroquine (CQ). MG132 is a reversible proteasome inhibitor. CQ is an autophagy inhibitor that can inhibit degradation through the lysosomal pathway. The result indicated that GMFB all R is degraded by proteasomal degradation systems (Fig. 4 D-F). Next, we overexpressed GMFB-wt with SUMO1 or SENP1 to enhanced GMFB SUMOylation or de SUMOylation in 293 cells, then measured the protein level post-treat with MG132 or CHX. As expected, GMFB is decreased when SENP1 overexpression (Fig. 4 G-H). SUMOylation protects GMFB from proteasome-mediated degradation. To further study the role of different “K” SUMOylation of GMFB in the stability of protein, we repeat the experiment with 293 cells transfected with GMFB wt-FLAG or GMFB K to R series plasmid and the MG132 or CHX assay was performed. We found that the mutation of K17, K38, K40, K74, K119, K137 did not affect protein stability (Fig. 4 I, J). These findings indicated that GMFB SUMOylation regulates protein stability.

**Figure 4.**
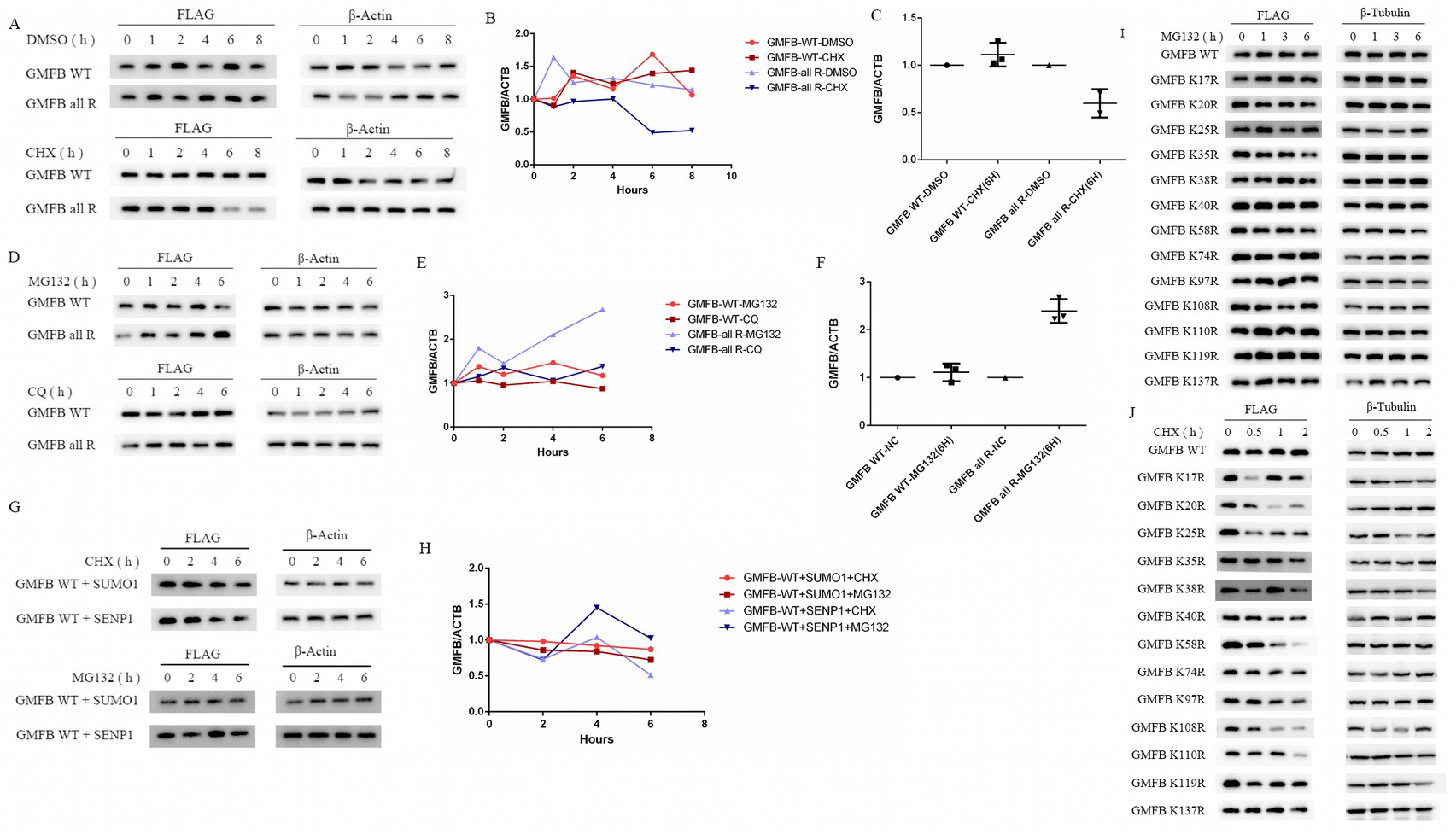
GMFB SUMOylation regulates protein stability (A) Western blot of total cell lysates from 293 cells transfected with GMFB wt-FLAG or GMFB all R-FLAG and treated with DMSO, CHX. Blot was probed using an anti-FLAG antibody. A representative blot is shown. (B) The graph shows the percentage of FLAG signals normalized to ACTB at the indicated time points in wild-type and all R mutant. (C) Quantification of normalized levels of GMFB wt-FLAG and GMFB all R-FLAG from western blots from 3 independent experiments for each treatment. (D) Western blot of total cell lysates from 293 cells transfected with GMFB wt-FLAG or GMFB all R-FLAG and treated with MG132 or CQ. Blot was probed using an anti-FLAG antibody. (E) The graph shows the percentage of FLAG signals normalized to ACTB at the indicated time points in wild-type and all R mutant. (F) Quantification of normalized levels of GMFB wt-FLAG and GMFB all R-FLAG from western blots from 3 independent experiments for each treatment. (G) Western blot of total cell lysates from 293 cells transfected with GMFB wt-FLAG with SUMO1 or SENP1 and treated with MG132 or CHX. Blot was probed using an anti-FLAG antibody. A representative blot is shown. (H) The graph shows the percentage of FLAG signals normalized to ACTB at the indicated time points in wild-type GMFB with SUMO1 or SENP1. Western blot of total cell lysates from 293 cells transfected with GMFB wt-FLAG or GMFB K to R series plasmid and treated with MG132(I), CHX(J). Blot was probed using an anti-FLAG antibody.

### SUMOylated GMFB translocated to nucleus and plasma membrane

SUMOylation is also known to regulate protein translocation [28]. Thus, we detected the cellular localization of GMFB in ARPE-19 cells transfected with sentrin-specific protease 1-targeting small interfering RNA (siSENP1). The result of immunofluorescence staining demonstrated that no observed nuclear translocation of GMFB, but GMFB seems to cluster around the membrane areas under high levels of global SUMOylation (Fig 5A). Next, we construct a fusion protein plasmid, FLAG-SUMO1(aa1-96)-GMFB, to mimic the dynamic balance of SUMOylated GMFB (32–34). The western blotting also confirmed FLAG-SUMO1-GMFB fusion proteins can cleaved by SENP1(Fig. 5B). The quantitative analysis of SUMOylated GMFB/free GMFB ratio in nuclear, cytosol and membrane fractionation in ARPE-19 cell line transfected with FLAG-SUMO1-GMFB plasmid indicated that SUMOylated GMFB may translocate to nucleus and membrane (Fig. 5 E-F). We also noticed that there are multiple bands of the SUMOylated GMFB in the nucleus, indicated other post-translational modifications are present in the nucleus.

**Figure 5.**
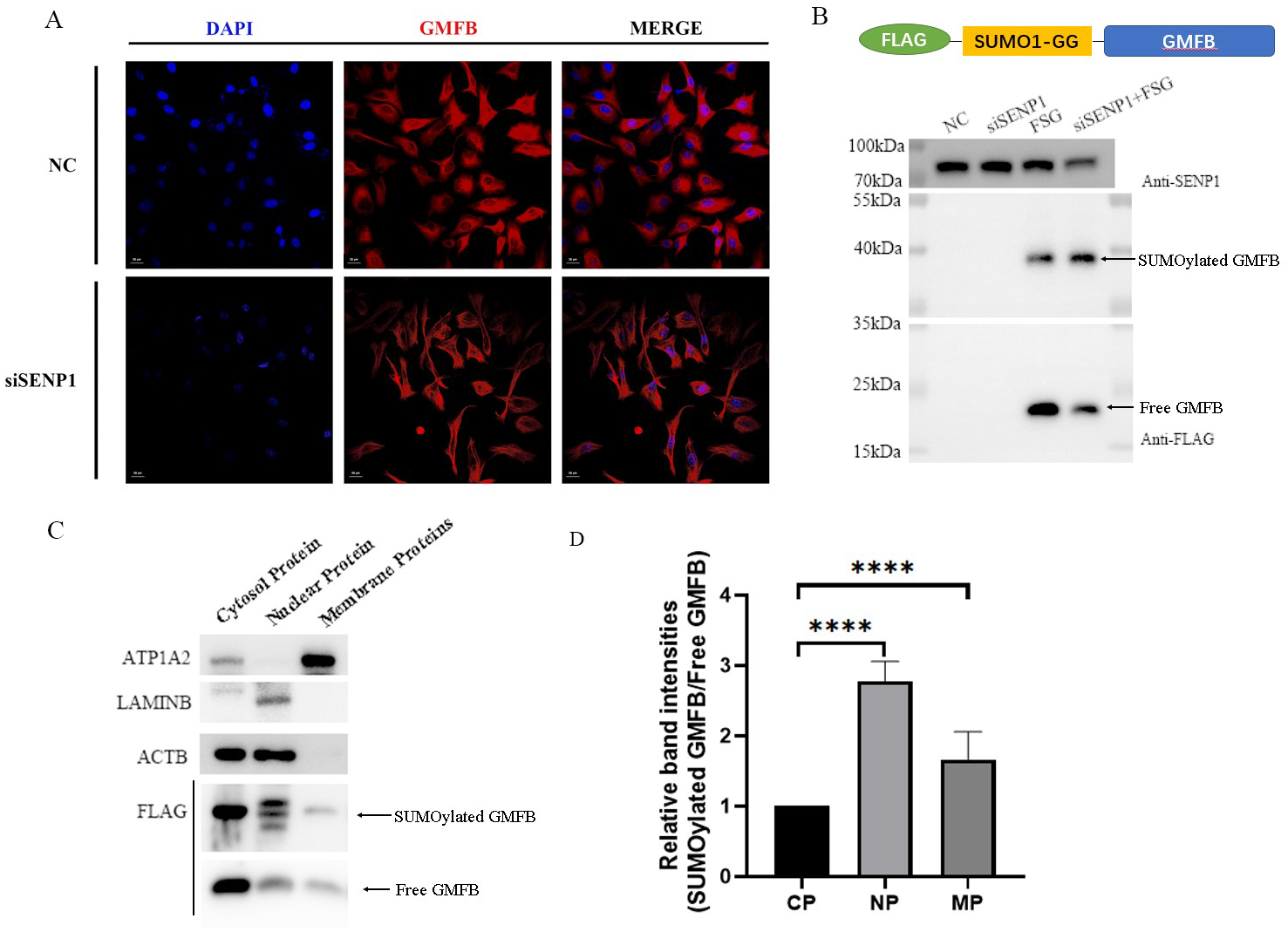
SUMOylated GMFB may be translocated to the nucleus and plasma membrane. (A) ARPE-19 cells were transfected siSENP1 in a 48-wells plate for 48 and stained by immunofluorescence with anti-GMFB antibodies and DAPI. (B) 293 cells were transfected with expression vectors for FLAG-SUMO1-GMFB and siSENP1, and immunoblot with antibody to FLAG and SENP1. (C) Nuclear, cytosol and membrane fractionation of GMFB in 293 cell line after transfected with FLAG-SUMO1-GMFB plasmid. ATP1A2, Lamin B and ACTB serve as membrane, nuclear and cytoplasmic markers, respectively. (D) The quantitative analysis of the gray density of Western blotting bands from GMFB of nuclear in 293.

### GMFB regulated the multiple pathways through SUMOylated modification

To unveil the molecular function of SUMOylated GMFB in RPE cells, we overexpressed empty vector (EV), GMFB-wt, GMFB-all R and GMFB K58R in ARPE-19 cells, then performed whole transcriptomic sequencing and comparative analysis. Hundreds of differentially expressed genes (DEGs) were identified among NC, GMFB wt, GMFB-all R and GMFB K58R. As shown in the volcano plot 369 genes were upregulated and 124 were downregulated, when overexpression GMFB wt (Fig. 2A). Among them, 66 upregulated DEGs were suppressed and 16 downregulated DEGs were increased by GMFB all R (Fig. 6B). Through analysis of PPI sub-networks, scores of differentially expressed genes (DEGs) were identified to interact with other proteins and found lots of genes associated with the cytokine-cytokine receptor (Fig. 6C). The top twenty most abundant KEGG pathways indicated that the SUMOylated GMFB upregulated multiple pathways, including the NOD-like receptor signaling pathway, TNF signaling pathway, RIG-I-like receptor signaling pathway, and NF-kappa B signaling pathway (Fig 6D). Furthermore, the heatmap of DEGs shows GMFB up-regulated a large number of genes. In contrast, GMFB all R resulted in down-regulation of partial genes. Similar results were obtained in the K58R group, but the effect was slightly weaker than GMFB all R group (Fig. 6E). Figure 7 showed that cytokine and cytokine receptor interaction up-regulated by the GMFB-wt and downregulated by GMFB-all. To validate the RNA-seq results in vitro and in vivo, the DEGs especially cytokine-cytokine receptors were selected for quantitation in the ARPE-19 cells transfected with EV, GMFB-WT and GMFB ALL R and K58R samples and the RBCC of wt and GMFB knock out rats, the results are consistent with RNA-seq (Fig. 6F-H).

**Figure 6.**
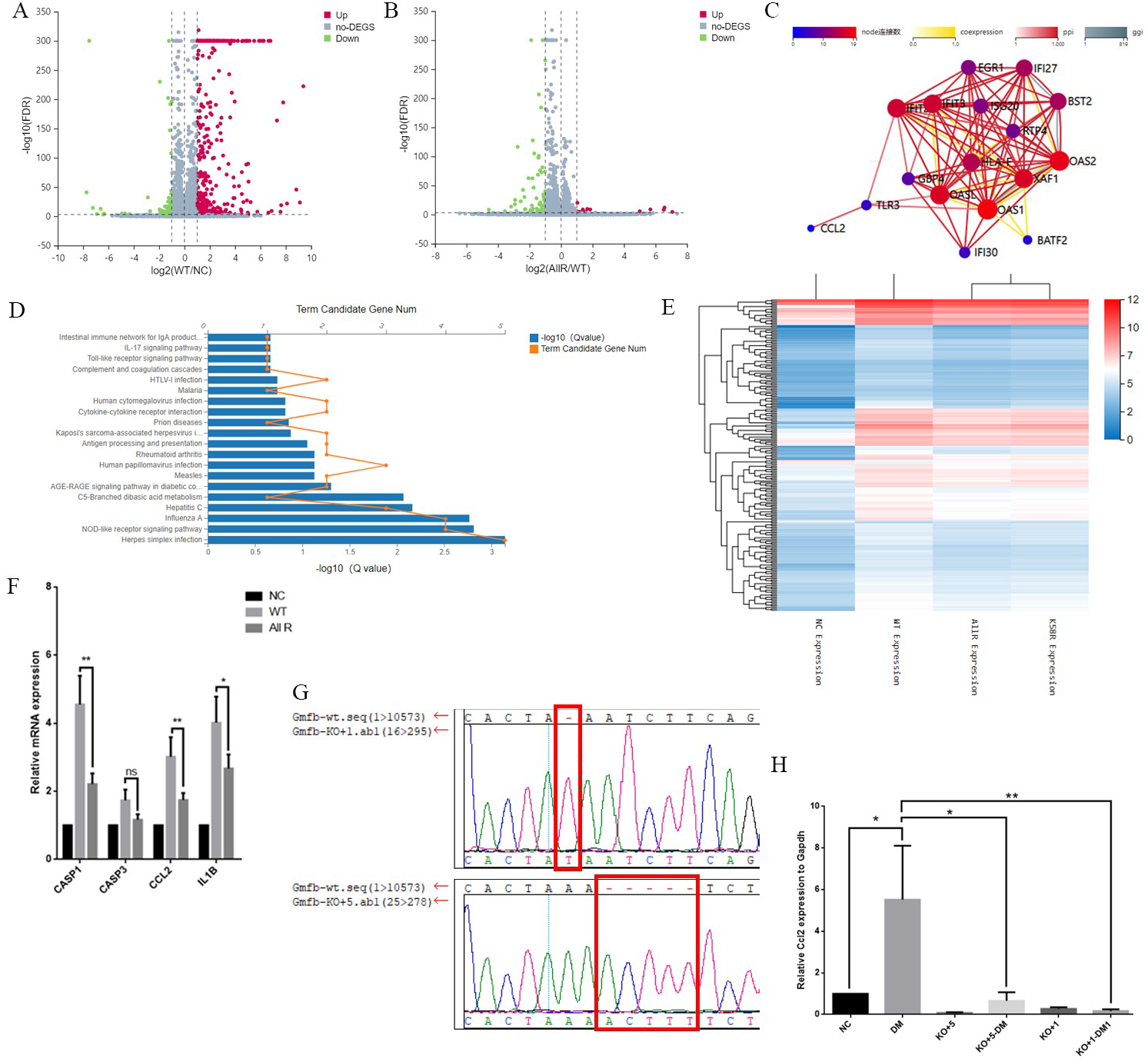
GMFB regulated the multiple pathways through SUMOylated modification. Volcano plot pictures showing the differentially regulated genes in both comparisons. (A) WT versus NC, (B) All R versus WT. The volcano plot was used as a filter to view the differentially expressed genes. A volcano plot shows the log2(Fold Change) in the x-axis against the –log10(FDR) in the y-axis. upregulated in red and downregulated in green. (C) The PPI network of DEG in the comparison of All R versus WT. (D) The top twenty most abundant KEGG pathways enrichment analysis for the ARPE-19 cells transfected with GMFB-wt and GMFB all R. (E) The heat map of the expression level of genes differentially regulated genes in the comparison of NC, wt, all R and K58R. (F) Several gene expression was measured on ARPE-19 cells transfected with GMFB-wt and GMFB all R by qPCR relative to ACTB. (G) Validation of Gmfb knockout rats by genomic DNA PCR and sequencing. (H) Ccl2 expression was measured on RBCC of NC, DM, KO+1, KO+1-DM, KO+5 and KO+5-DM rats by qPCR relative to Gapdh. Bars represent mean ± SEM (n=7). **P<0.01; ***P<0.001. Bars represent mean ± SEM. **P<0.01; ***P<0.001; ****P<0.0001.

**Figure 7.**
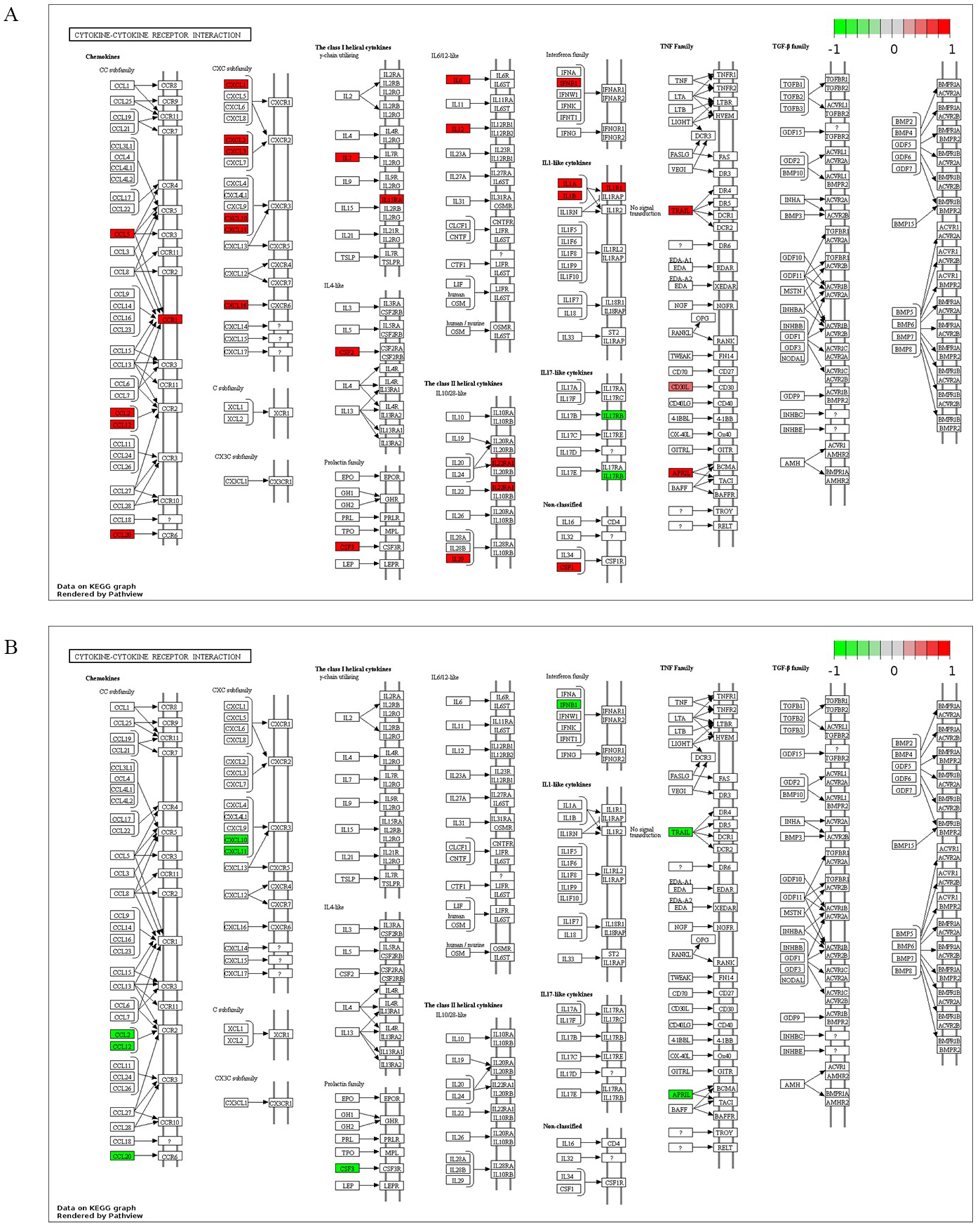
The KEGG pathway was constructed by the Kyoto Encyclopedia of Genes and Genomes (KEGG) pathway database. Cytokine and cytokine interaction up-regulated by the GMFB-wt (A) and down-regulated by GMFB-all (B).

## Discussion

GMFB is highly evolutionary conserved between yeast and mammals and regulates inflammation in the central nervous system (CNS)(4,35). In mouse primary astrocytes, overexpression GMFB leads to upregulation of nuclear factor-κB (NF-κB) and a significant increase of granulocyte-macrophage colony-stimulating factor (GM-CSF), a pro-inflammatory cytokine, secretion(36). The SUMOylation also has been considered to play a role in inflammatory processes. Previous research showed that BMSCs overexpressing SUMO are more tolerant to hypoxia conditions, and have stronger repair ability for damaged chondrocytes in vitro and for articular cartilage injury model in rats, and are excellent seed cells for repairing articular cartilage(37).

Previous work has shown that hypoxia enhances protein SUMO-1 expression and modification, inhibits the expression of the SENP1(38–41). The oxidative and ER stress also increase global SUMOylation(42). In our study, we identified the expression of GMFB and SUMO1 was enhanced under the early stage of hypoxia/high glucose or oxidative stress in RPE cells. Our conclusions are consistent with the former research. Therefore, the role of SUMOylated GMFB in RPE cells under oxidative stress and inflammation has been examined.

The human genome encodes four homology SUMO isoforms, which are SUMO1, SUMO2, SUMO3 and SUMO4(43). SUMO4 expression mainly in adults and embryonic kidneys(44). As for SUMOylation, SUMO1 is mainly conjugated to proteins as the monomeric form, SUMO2/3 is able to form polymeric chains(18,45). SUMO1 cannot form chains, but it can act as a chain terminator(46). The sodium iodate-inducible model was established to represent oxidative stress and inflammation in the RPE layer. The expression patterns of SUMOs in RBCC indicated SUMO1 is more sensitive at the initial stage of the reaction(29). Therefore, we hypothesized and confirmed that GMFB is covalently modified by SUMO1 at only one SUMOylated site.

Subsequently, we affirmed that there is no specific SUMOylated site in GMFB, which means that the SUMOylated site in GMFB can shift. Moreover, we identified 4 major SUMOylated sites, 20k, 35k, 58k and 97k, even without the consensus sequence flanking lysine, the SUMOylation can still occur(47).

Based on the current research state, SUMOylation enhances the protein stability via binding to lysine residues that otherwise would be ubiquitinated (48,49). Our functional studies also reveal that the stability of GMFB protein is regulated by SUMOylation. And the stability of GMFB in RPE cells is critical in maintaining the function of GMFB.

GMFB is primarily an intracellular protein because of lacking signal peptide sequence(50). Our results confirm that the SUMOylated GMFB translocates to membrane and nuclear. A previous study showed that GMFB is expressed on the cell surface of astrocytes and thymus epithelial cell lines support our result(51,52). There has been reports showing GMFB can be secreted under certain conditions(53,54). In the rat brain, GMFB has been found to be increased by 7-fold in the wound cavity after aspiration lesion after rat brain injury and leading to glial proliferation(54).

RNA-Sequencing data indicates that overexpression GMFB is characterized by increased expression of genes that are related to the inflammatory responses (OAS1, DHX58, CASP1, TNFSF13, RASGRP3, NFKBIA, HLA-A, MYD88, TNFAIP3, STAT2, IL1A, IFITM2, IL15RA, IFITM1, IFI30, GBP4, TLR3, and IL1B). De-SUMOylated GMFB is also characterized by decreased expression of genes that are related to the inflammatory system (OAS1, TNFSF13, IFI30, GBP4, TLR3, CCL2, SERPING1, HLA-F, and OAS2). Moreover, the construction of the protein-protein interaction (PPI) network, recognition of hub genes and cluster analysis provides insight into the molecular mechanisms of SUMOylated GMFB. High GMFB expression level upregulating NOD-like receptor signaling pathway, TNF signaling pathway, RIG-I-like receptor signaling pathway, cytokine-cytokine receptor interaction and NF-kappa B signaling pathway, and these pathways were down-regulated by de-SUMOylated GMFB. Therefore, SUMOylated GMFB also plays an important role in inflammatory responses.

NOD-like receptor signaling pathway participates in several ocular diseases such as DR(55). A previous study has shown that sustained activity of the NOD-like receptor signaling pathway can activate the pro-inflammatory cytokines IL-1β and IL-18 in type 2 diabetic patients(56). The secretion of IL-1b from RPE cells also via the NLRP3 Inflammasome(57). Furthermore, the findings support that the SUMOylated GMFB participates in DR pathogenesis.

The C-C motif chemokine 2 (CCL2), also called MCP-1, is a member of the CC chemokine family that plays a vital role in DR(58). It is generally known that expression levels of CCL2 were significantly associated with the clinical stages of DR, and considered as therapeutic targets in DR(59). The retinal degeneration induces upregulation of the secretion of inflammatory chemokines such as CCL2 via RPE cells, Muller cells and microglia in DR (60). Thus, the RT-qPCR assay was performed to verify the role of SUMOylated GMFB on CCL2 expression in vitro and in vivo. However, the specific mechanisms underlying SUMOylated GMFB in DR require further study.

Although there are important discoveries revealed by these studies, there are also limitations. First, it’s difficult to detect endogenous SUMOylated GMFB due to dynamic posttranslational modification process; Second, we failed to detect SUMOylated site by mass spectrometry because of technical limitations ; Third, further work needs to be performed to investigate the role of SUMOylated GMFB in many ophthalmic diseases.

In summary, the present study demonstrates that SUMOylated GMFB is involved in a variety of inflammatory signaling pathways and plays crucial roles in ocular inflammatory disorders such as diabetic retinopathy. These results help to clarify the association between SUMOylation, GMFB and inflammation, suggesting a new target for the treatment of RPE-related disorders like age-related macular degeneration (AMD) and diabetic retinopathy (DR).

## Experimental procedures

### Cell lines

ARPE-19 and HEK293T cell purchased from ATCC were used in this study. All cells were maintained in Dulbecco's Modified Eagle Medium (DMEM) supplemented with 10% FBS (Gibco, Thermo Fisher Scientific, Spain) at 37 °C and 5% CO2. and cultured.

### Plasmids

To construct pCMV-flag-WT-GMFB, pCMV-His-SUMO1, pCMV-UBC9, the DNA fragments were amplified by PCR and subsequently cloned into the vector pCMV. Site-directed mutagenesis was used to make amino acid changes, using pCMV-flag-WT-GMFB as templates.

### Cell transfection

HEK293T and ARPE-19 cells were transfected using Lipofectamine 2000 (Invitrogen, USA), according to the manufacturer's instructions. Plasmid and Lipofectamine 2000 was diluted in FBS free DMEM(Gibco) medium and incubated for 5 min, separately. Then mixed together, and incubated for 10-15 min at room temperature (RT) to form the DNA-Lipofectamine complexes. Finally added to the culture plates.

### CHX, CQ and MG132 assay

To examine protein stability, cells were seeded into 24-well plates, cultured for 24 h, and transfected with indicated plasmids. After treatment with CHX (20 μg/mL), CQ(50 μM), or MG132(20 uM) for indicated times, cells were then harvested, and subjected to western blotting.

### STZ-induced diabetes model

Male Sprague Dawley rat (150 g) fasted overnight before given intraperitoneal injections of 50 mg/kg STZ dissolved in a 10mM sodium citrate buffer solution, pH of 4.5. Animals with blood glucose levels over 300 mg/dl were considered diabetic.

### siRNA knockdown

For knock-down experiments, ARPE-19 cells were seeded at 50%–60% confluence in six‐well tissue culture plate and grown overnight. Cells were transfected with double‐stranded inhibitory RNA oligonucleotides using Lipofectamine 3000 (Invitrogen, USA) according to the manufacturer’s instructions. Western blot analyses were carried out to confirm the specific inhibitory activity.

### Immunoprecipitation and Western blot analysis

Immunoprecipitation and western blotting assays were performed as previously described. Cells were transfected with plasmid for 48 h and washed twice in ice-cold PBS, lysed in an immunoprecipitation (IP) buffer for 30 min. The lysates were centrifuged at 12,000 g for 20 min to collect supernatants, immunoprecipitated with specific beads, rotated at 4°C for 4 h. Immunoprecipitates were washed three times with IP buffer before loaded and analyzed via SDS-PAGE separation and immunoblotting.

### Immunofluorescence

The slices of cells or tissues were washed 3 times in PBS then fixed with 4% polyformaldehyde (Solarbio) at room temperature for 1 h. cells were incubated with 0.5% Triton X-100 (Solarbio) for 5 min, followed by permeabilized in 0.1% Triton for 15 min. Afterwards, blocked with 1% bovine serum albumin(BSA) for 1h with the following primary antibodies: GMFB(1:100), SUMO1(1:50) overnight at 4°C. Afterwards, cells were incubated with conjugated secondary antibody (1: 200) for 1 h following three 5-min washes in PBS. The nucleus was stained by DAPI. Fluorescence signals were observed under confocal microscopy.

### RNA-Seq and KEGG pathway analysis

The illumina sequencing, Gene ontology (GO), pathway enrichment analysis, construction of the protein–protein interaction (PPI) network (PIN), recognition of hub genes and cluster analysis were completed by Beijing Genomics institution.

### quantitative real-time PCR

Total RNA was extracted using the TRIzol reagent (Takara, Dalian, China) according to the manufacturer’s instructions, then reverse transcribed to cDNA using PrimeScript RT polymerase (Takara, Dalian, China). SYBR Green Master Mix (Tiangen Biotech, China) on a LightCycler 96 Detection System (Roche) was used for RT-qPCR. Primers used in this study are listed in Table 2. Data were analyzed using the 2-ΔΔCq method.

**Table 2.**
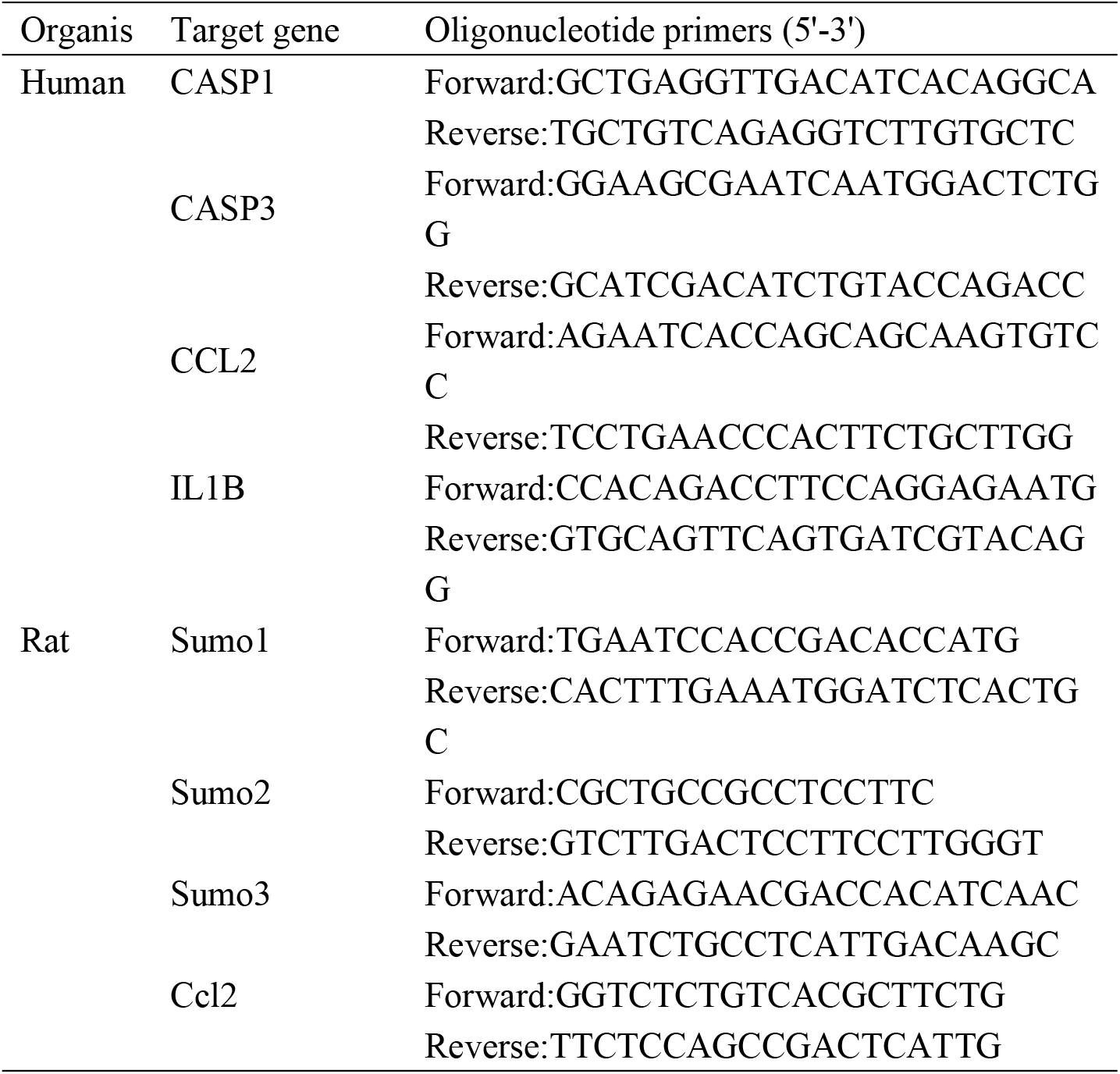
Primers used to amplify human CASP1, CASP3, CCL2 and IL1B, and rat Sumo1, Sumo2, Sumo3 and Ccl2.

## Author contributions

Wan Sun performed the experiments, analyzed the data and wrote the main manuscript text; Lixia Lu, Guo-Tong Xu and Jian Huang conceived the experiments, analyzed the data and revised the manuscript. Juan Wang, Jieping Zhang, Furong Gao, Qingjian Ou, Haibin Tian, Caixia Jin, Jingying Xu, and Jingfa Zhang participated in analysis and discussion. All authors reviewed the manuscript and had final approval for the submitted version.

## Funding and additional information

We thank all members of our laboratory for many helpful discussions. This work was supported by grants obtained from the Ministry of Science and Technology of China (2017YFA0104100, 2015CB964601, 2016YFA0101302), the National Natural Science Foundation (81670867, 81372071, 81770942) and the Shanghai Municipal Commission of Health and Family Planning project (201640229), and the Shanghai Science and Technology Committee Grant (17ZR1431300), as well as a grant from Shanghai East Hospital (ZJ2014-ZD-002).

